# Parthenogenesis as a solution to hybrid sterility: the mechanistic basis of meiotic distortions in clonal and sterile hybrids

**DOI:** 10.1101/663112

**Authors:** Dmitrij Dedukh, Zuzana Majtánová, Anatolie Marta, Martin Pšenička, Jan Kotusz, Jiří Klíma, Dorota Juchno, Alicja Boron, Karel Janko

## Abstract

Formation of species generally occurs in a continuum from potentially intermixing populations to independent entities isolated from other species by pre- and postzygotic barriers. Especially the establishment of hybrid sterility (HS) is a hallmark of speciation, which usually emerges at different rates between hybrid sexes. However, although HS is frequently observed, the underlying molecular mechanisms remain poorly understood. Here we report that speciation proceeds through a previously unnoticed stage at which gene flow is completely interrupted on side of both hybrid’s sexes, although only male hybrids are sterile, while female fertility is rescued due to a particular gametogenetic deviation leading to the formation of clonal gametes. Specifically, analysis of gametogenetic pathways in hybrids between fish species Cobitis elongatoides and C. taenia revealed that male HS resulted from extensive asynapses and crossover reduction among elongatoides-taenia chromosomal pairs followed by apoptosis. By contrast, hybrid females exhibited premeiotic genome endoreplication which ensured proper formation of bivalents between identical chromosomal copies. This deviation ultimately restored fertility in females but since it simultaneously leads to the production of unreduced clonal gametes, it restricts interspecific gene flow thereby directly contributing to speciation. In conclusion, our data demonstrate that the emergence of asexuality may remedy HS in a sex-specific manner and is intermingled with the speciation process. Although gametogenetic mechanisms employed by asexual animals and plants have rarely been scrutinized, available evidence suggests that premeiotic endoreplication is relatively widespread. This suggests that observed link between HS and clonality may have general validity in taxa able of asexual reproduction.

**Author’s summary:** Species are fundamental evolutionary units that presumably evolve in a continuum from potentially intermixing populations to independent entities isolated from other species by pre- and postzygotic barriers. Especially the establishment of hybrid sterility (HS) is a hallmark of speciation, which usually emerges at different rates between hybrid sexes. However, although HS is frequently observed, the underlying molecular mechanisms remain poorly understood. Here we report the existence of a previously unnoticed stage of speciation at which gene flow is completely interrupted, although only male hybrids are sterile, while female fertility is rescued due to a particular gametogenetic deviation leading to formation of clonal gametes. Specifically, HS resulted from extensive asynapses in male gonads, but in females the hybridization provoked premeiotic endoreplication which rescued chromosome pairing and fertility. Simultaneously, this meiotic deviation caused clonal transmission of maternal genome, thereby effectively restricting the interspeficic gene flow. Our results emphasize that emergence of clonality is a type of hybrid incompatibility that is intermingled with the formation of biological species and may remedy hybrid sterility in a sex-specific manner.

## Introduction

Molecular machinery controlling the production of reduced gametes is highly conserved [1]. Nevertheless gametogenesis has been repeatedly modified during the course of evolution, giving rise to many taxa that employ a wide range of cytogenetic mechanisms to form more or less unreduced gametes, ranging from completely ameiotic processes (apomixis) to those with retained meiosis and even recombination (automixis); reviewed in [2,3]. For example, clonal gametes may be produced despite normal meiotic divisions when chromosomes undergo premeiotic endoreplication and subsequent pairing of identical copies [4]. Although these so-called asexual organisms are intensively studied by evolutionary biologists, surprisingly little is known about the mechanisms that initiate the switch from sexual to asexual reproduction. Many asexual organisms appear to have formed as a result of hybridization since they combine genomes of two or more sexual species. Although the reasons that deviation from sexual reproduction appears strongly connected to interspecific hybridization remain unknown, several theoretical frameworks have been proposed to explain this link. Moritz et al. [5] assumed that unreduced gametes may arise in a hybrid when hybridizing species accumulated sufficient amount of intergenic incompatibilities to disrupt the cellular regulation of sexual reproduction, yet not enough to compromise hybrid’s fertility. Alternatively, De Storme and Mason [6] accentuated the role of decreased sequence homology among chromosomes in preventing their pairing and segregation, while Carman [7] hypothesized that hampered cross-talk between diverged regulatory programs brought together by hybridization may cause the switch to asexual reproduction.

Although empirical support for any of these hypotheses is frustratingly scarce, they all accentuate the importance of increased divergence between hybridizing species. Over a century ago, Ernst [8] proposed that the type of a hybrids’ reproduction depends on divergence between its parental species following a continuum from sexual hybrids between close relatives to obligate asexual hybrids between distant species. This hypothesis was recently corroborated by recent analysis of speciation in spined loaches (*Cobitis elongatoides* and *C. taenia*) [9]. In the previous study we found that initial stages of species divergence were accompanied by massive gene flow, thereby evidencing the existence of hybrids with efficient recombination and segregation, but at later stages, gene flow ceased as all hybrids produced by contemporary species are either clonally reproducing (females) or sterile (males). To further test the correlation between interparental divergence and reproductive strategy of hybrids, we compared published data on hybrids among ray-finned fishes and found that genetic divergences of species pairs producing asexually reproducing hybrids are higher than of those producing sexual hybrids, but lower than of those producing only sterile hybrids [9].

It is important to note that clonal reproduction in hybrids efficiently restricts interspecific gene exchange [9,10] and hence, it may isolate nascent species before the emergence of other classical forms of reproductive incompatibility [9]. Thus, a century after Ernst, it became clear that theories relating the initiation of asexuality to hybridization have many analogies to speciation models assuming the accumulation of postzygotic reproductive incompatibilities (RIM). Indeed, the hybridization-asexuality theories and the Bateson-Dobzhansky-Muller (BDM) class of speciation models both relate the establishment of the central trait (i.e. asexuality and RIM, respectively) to the genetic distance between hybridizing species. Because intergenomic incompatibilities presumably accumulate stochastically, both concepts also predict that the central trait may be achieved via a wide spectrum of molecular and cytological pathways (see e.g. [2,11]).

An important feature common to both RIM accumulation and hybrid asexuality is their tendency to emerge in a sex specific manner. Early stages of speciation are often characterized by a prominent decrease in fertility of one hybrid sex only, which may relate to the sex determination system [12], but the basis of such asymmetry still remains elusive. For example, although both hybrid sexes ultimately achieve sterility at advanced stages of speciation, it is unclear whether males and females proceed by the same or by different routes (e.g. [13,14]). Similarly, asexually reproducing taxa are usually called “unisexual” or “all-female” [15] but it is generally unclear whether apparent female-bias results from sex-specificity of gametogenetic aberrations or whether asexual males are generally ignored as they may not generate progeny on their own apart from some exceptions like androgenesis [16]. Interestingly, when hybrid brothers of known asexual females have been sought, they are generally documented as sterile or absent (e.g. [17,18], but see e.g. [19] for exceptions).

In this study we reveal proximate gametogenetic aberrations leading to hybrid asexuality and sterility in the so-called *Cobitis taenia* hybrid complex (Teleostei). We focus on two species that diverged ca 9 Mya, *C. elongatoides* and *C. taenia* [9] but regularly hybridize and produce viable diploid hybrids. Hereafter we denote haploid genomes as “E” and “T”, so that e.g. pure *C. elongatoides* specimens are called “EE” and diploid hybrids “ET”. Reproductive experiments and histological observations showed that *C. elongatoides-taenia* hybrid females (both diploid and triploid, naturally occurring as well as artificial F1 strains) are fertile and reproduce clonally by sperm-dependent parthenogenesis, known as gynogenesis (i.e. they produce unreduced eggs but require sperm to activate cellular division; [9,20–23]. Occasional fusion of sperm with hybrid egg pronuclei results in triploid progeny with EET or ETT genomic constitution. In contrast to their clonal sisters, the male hybrids have drastically reduced reproductive capabilities and appear incapable either of fertilizing the normal haploid eggs or of triggering the development of the clonal eggs [21,23,24]. Any interspecific gene flow is thus prevented by clonality (females) or sterility (males) of contemporary hybrids. Fig 1A demonstrates the aforementioned reproductive interactions among genomotypes of the *C. taenia* complex.

**Fig 1.**
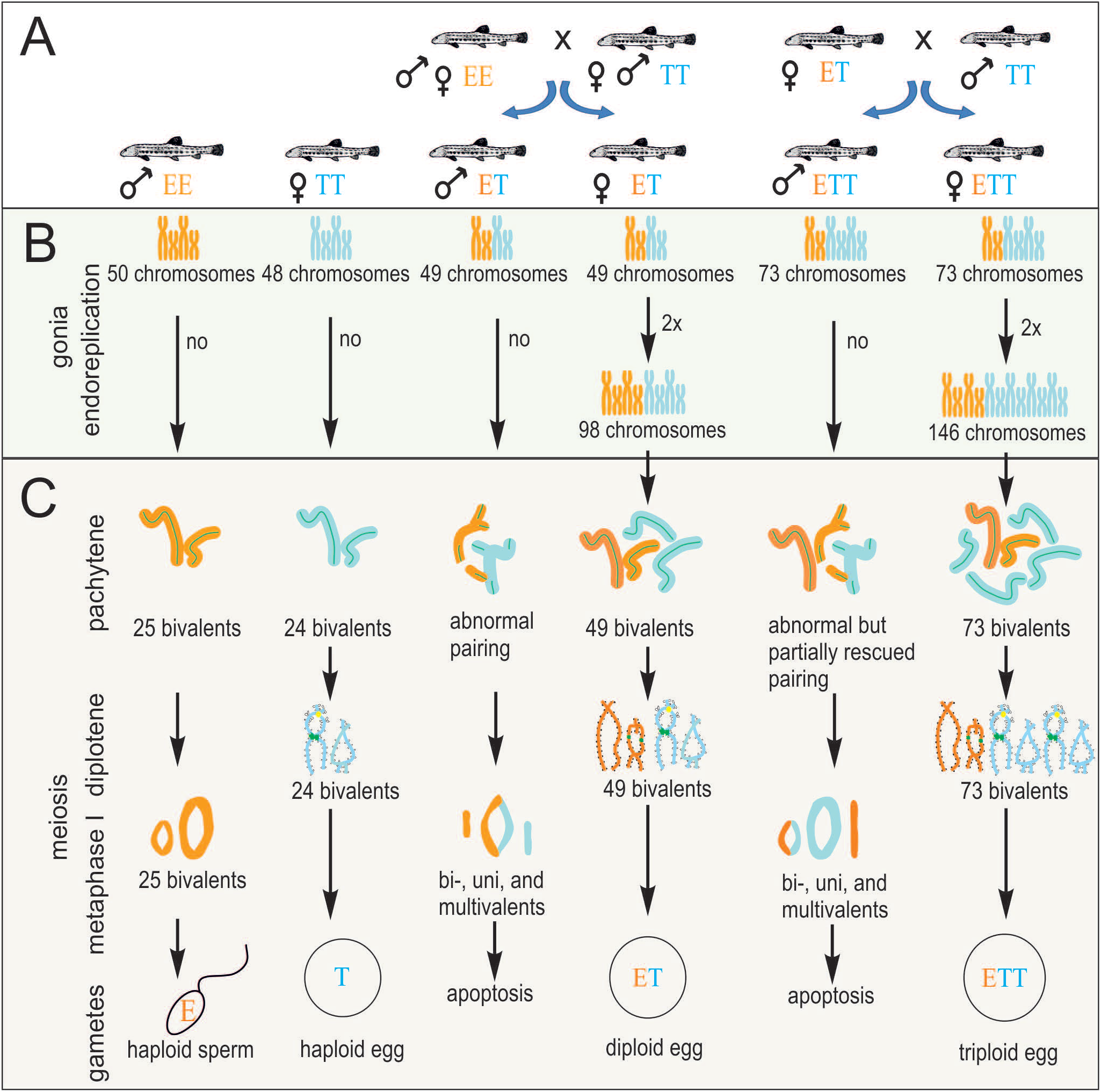
Schematic overview of gametogenetic pathways and aberrations in parental species and hybrids of both sexes. (A) scheme of reproductive interactions among studied genomotypes and the types of resulting progeny. (B) hypothetical gametogenetic pathways before meiosis. (C) observed stages of meiosis. Note that for methodological reasons, we studied pachytene stage in both sexes, but diplotene and metaphase I stages could be observed only in females and males, respectively.

This study is the first to reveal the mechanistic basis of simultaneously arising sterility of hybrid males and asexuality of hybrid females including the effects of genome duplications and polyploidy on hybrid gametogenesis. We show that sex-specific cytogenetic mechanisms of parthenogenetic reproduction prevent the occurrence of sterility in hybrid females. Still, although parthenogenesis restores female fertility, it provides an efficient barrier to interspecific gene flow and thereby contributes to speciation as much as male sterility.

## Results

### Hybrid sterility and asexuality evolves in a sex specific manner

Mechanistically, reproductive incapacity of hybrid males is reflected by defective development of hybrid testes as evidenced by histological examination of diploid ET ([23], this study) and triploid ETT hybrid males (this study). The testes of sexual as well as hybrid males, contained properly developed Sertoli and Leydig cells but germinal cells in cysts of hybrid testes developed asynchronously as demonstrated in Fig 2 A, A’ and in detail in S1 Fig). Importantly, hybrid spermatogonia and spermatocytes at the beginning of prophase I (leptotene/pachytene) were properly developing and possessed typical organelles for such stages, such as mitochondria, Golgi apparatus, “nuage,” nucleus and nucleolus (Fig 2B, B’, 2C, 2C’). In metaphase I, the chromatin of all spermatocytes of sexual males was equatorially distributed, while spermatocytes of hybrids demonstrated irregular chromatin distribution; (Fig 2D, D’). After metaphase I, we usually observed only fragmented postmeiotic germ cells with numerous nuclear vesicles and multiple axonemes/flagella in hybrids (Fig 2 E, E’). In very rare cases, testes of hybrids contained some asynchronously developing cysts with a few typical spermatozoa, which nonetheless often contained multiple axonemes or flagella. The nuclei of those spermatozoa were generally larger (Fig 2 F, F’), which may indicate higher DNA content or abnormal chromatin packaging. Hybrid testes contained many degenerating germinal cells, which were more often found among spermatocytes than among spermatogonia and we documented frequent apoptotic processes there (for detailed sexual-hybrid comparison see S2 Fig).

**Fig 2.**
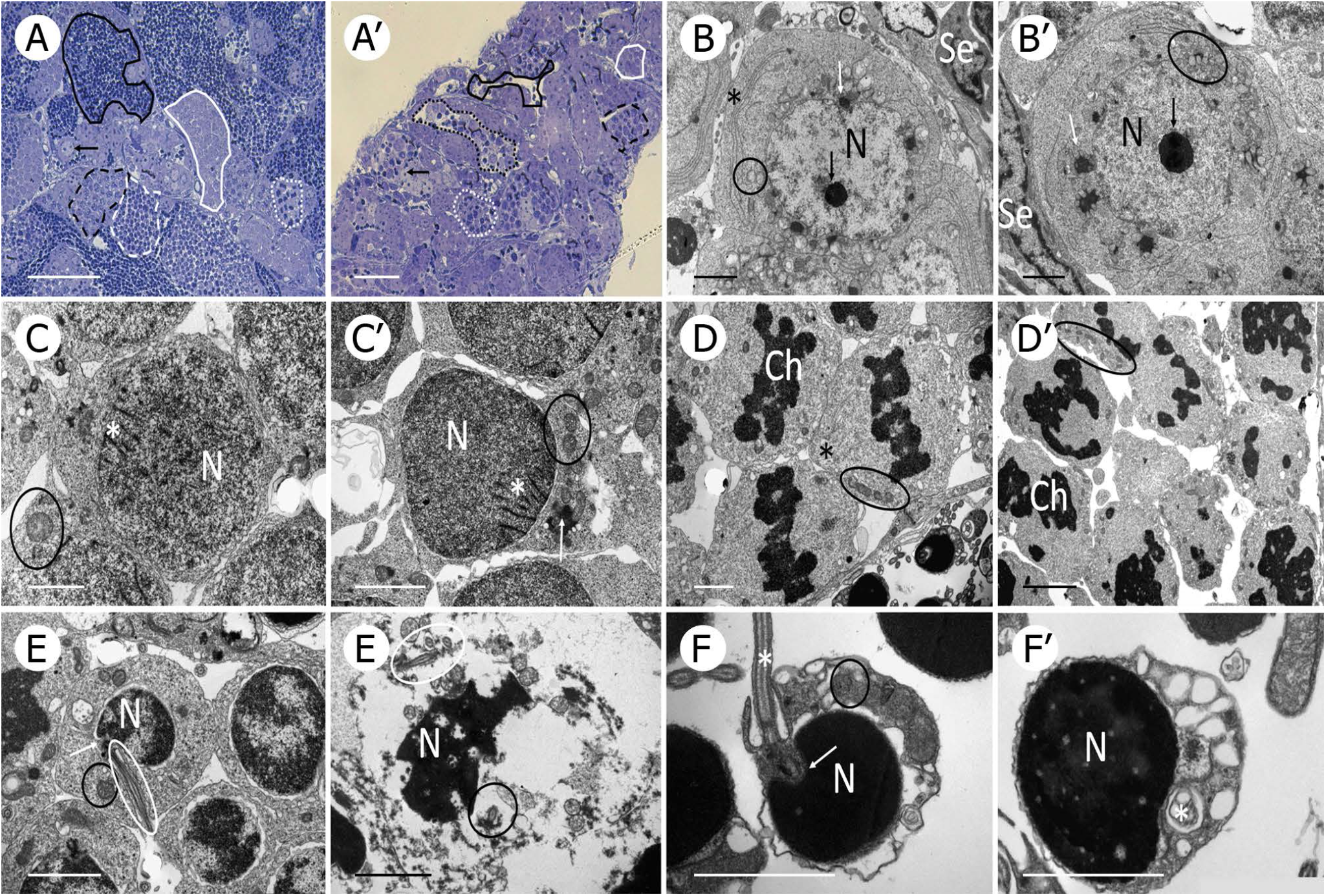
Comparison of spermatogenesis between male representatives of sexual diploid species (panels A, B, C, D, E, F) and hybrid genomotype (panels A’, B’, C’, D’, E’, F’). As the variation between two species and between two hybrid genomotypes is negligible, for simplicity we selected *C. elongatoides* and ETT male as representatives of both groups. (A-A’) semithin sections show spermatogonia (arrow), spermatocyte in zygotene/leptotene (white firm line) and pachytene (black dashed line) of prophase I, spermatocyte in metaphase I (white dotted line), spermatids (white dashed line) and spermatozoa (black firm line), while testes of hybrid displays defective development with only a few single spermatozoa and germ cells of one cyst at different stages (black dotted line); (B-B’) spermatogonia A with nucleus (N), nucleolus (black arrow), Golgi apparatus (asterisk), mitochondria (black circle), nuage (white arrow) and Sertoli cell (Se); (C-C’) spermatocyte in zygotene/leptotene of prophase I with nucleus (N), mitochondria (black circle), nuage (white arrow) and synaptonemal complexes (white asterisk); (D-D’) spermatocytes in metaphase I show compact chromatin (Ch) in equatorial position with spindle fibers (black asterisk) and mitochondria (black circle) (on the contrary, hybrid demonstrate irregular distribution of chromatin and no spindle fiber formation); (E-E’) spermatids in sexual diploid display nucleus (N), mitochondria (black circle), basal body (white arrow) and flagellum (white circle), while spermatids of hybrids are usually fragmentized and contain numerous axonemes/flagella; (F-F’) spermatozoa are composed of nucleus (N), mitochondria (black circle), basal body (white arrow), and flagellum (white asterisk) (hybrids demonstrated very rare occurrence of spermatozoa with generally bigger nucleus).

### Gametogenetic pathways leading to clonality or sterility differ by sex-specific premeiotic stage

Cytogenetic analysis identified basic differences between sexes in the development of germ cells and subsequent meiosis, which probably represent a proximate reason for asymmetrical evolution of hybrid asexuality and sterility. In both parental species, the nuclei spreads from oocytes at the diplotene stage showed, as expected, the same numbers of chromosomes as their somatic cells (i.e. 48 in *C. taenia* and 50 in *C. elongatoides*), which paired into bivalents whose numbers equaled exactly half of these counts (i.e. 24 and 25, respectively; Fig 3A, B). By contrast, the number of bivalents in diploid ET and triploid ETT diplotene nuclei always equaled the number of chromosomes in their somatic cells (i.e. 49 in ET and 73 in ETT; Fig 3C, D). Exclusively bivalents were observed in hybrid female oocytes with no evidence for uni- or multivalents.

**Fig 3.**
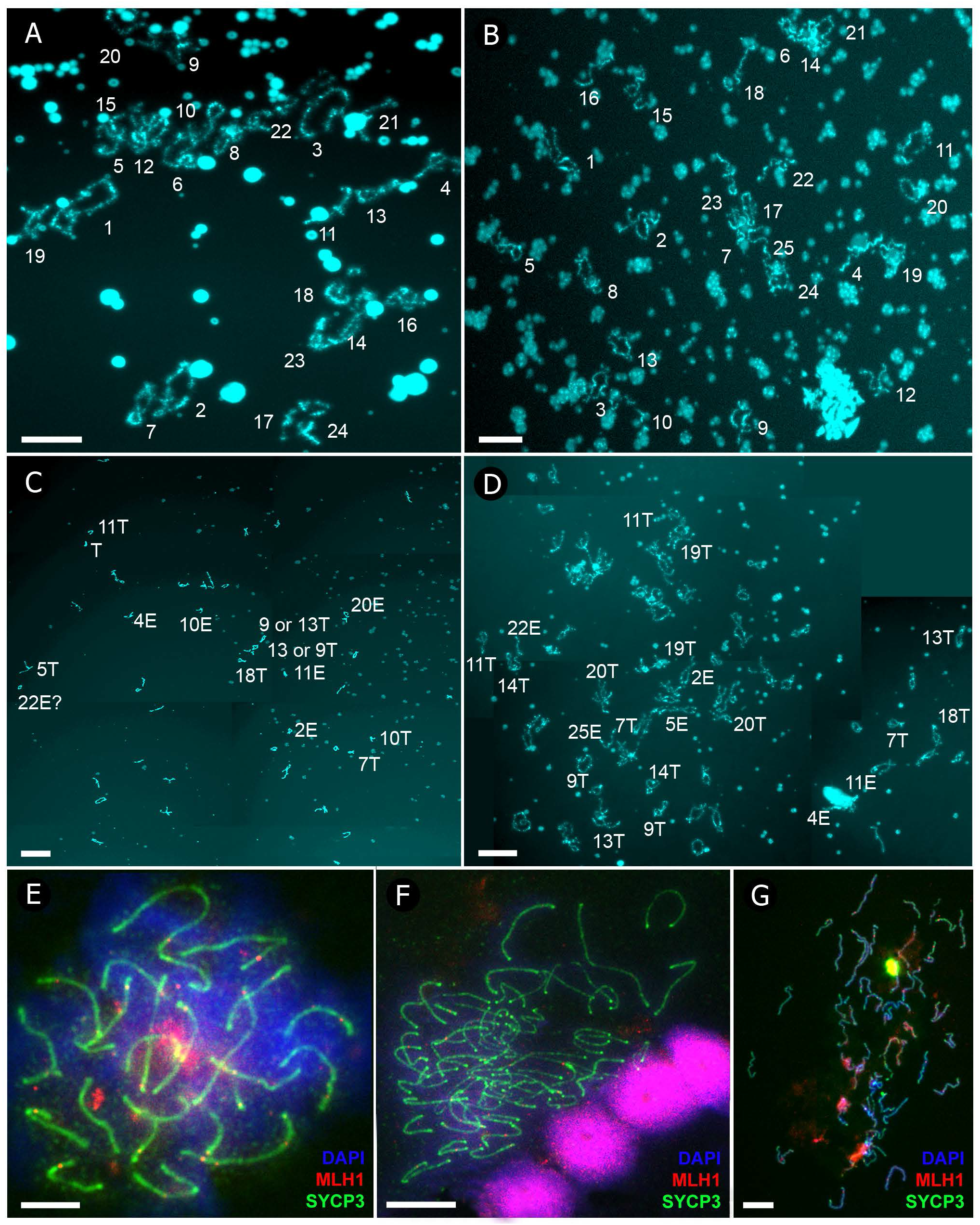
Female meiotic spreads at diplotene (A-D) and pachytene (E-G) stages. Panels A-D demonstrate the spreads of lampbrush chromosomes from diplotene oocytes of *C. taenia* with 24 bivalents (A), *C. elongatoides* with 25 bivalents (B), diploid ET hybrid with 49 bivalents (C) and triploid ETT hybrid with 73 bivalents (D). Lampbrush chromosomes are numbered numerically according to their size and morphology (see S2 Fig for detailed map of lampbrush chromosomes). Subscripts in italics indicate the distinguishable lampbrush chromosomes unequivocally corresponding to *C. elongatoides* “e” and *C. taenia* “t”, respectively. Scale bar = 50µm. Panels E-G demonstrate spread pachytene oocytes of *C. elongatoides* (E), diploid ET (F) and triploid ETT (G) hybrid females stained with DAPI (blue); synaptonemal complexes (SCs) were immunolabelled with antibodies against SYCP3 protein (green) and MLH1 protein (red). Scale bar = 10µm.

This contrasts with our findings for the hybrid ‘brothers’ of these asexual females. We analysed approximately 250 metaphase I nuclei in each male, and noticed that they always contained the same number of chromosomes as the somatic tissues, i.e. 50, 49 and 73 in EE, ET and ETT genomotypes, respectively Fig 4A, B, D. With the help of telomeric FISH probes we detected 25 and 24 bivalents in *C. elongatoides* and *C. taenia* males, respectively, but in hybrids, few chromosomes paired into bivalents while most formed uni- and multivalents. On average we observed ~ 34 univalents, 5 bivalents and 1 multivalent, among the 49 chromosomes of diploid ET males, while ETT triploids possessed a considerably higher proportion of bivalents with an average of of ~ 23 univalents, 12 bivalents and 8 multivalents among their 73 chromosomes. In any case, observed patterns strikingly contrast with those of females and suggests that premeiotic endoreplication is not a prevailing gametogenetic aberration in hybrid males.

**Fig 4.**
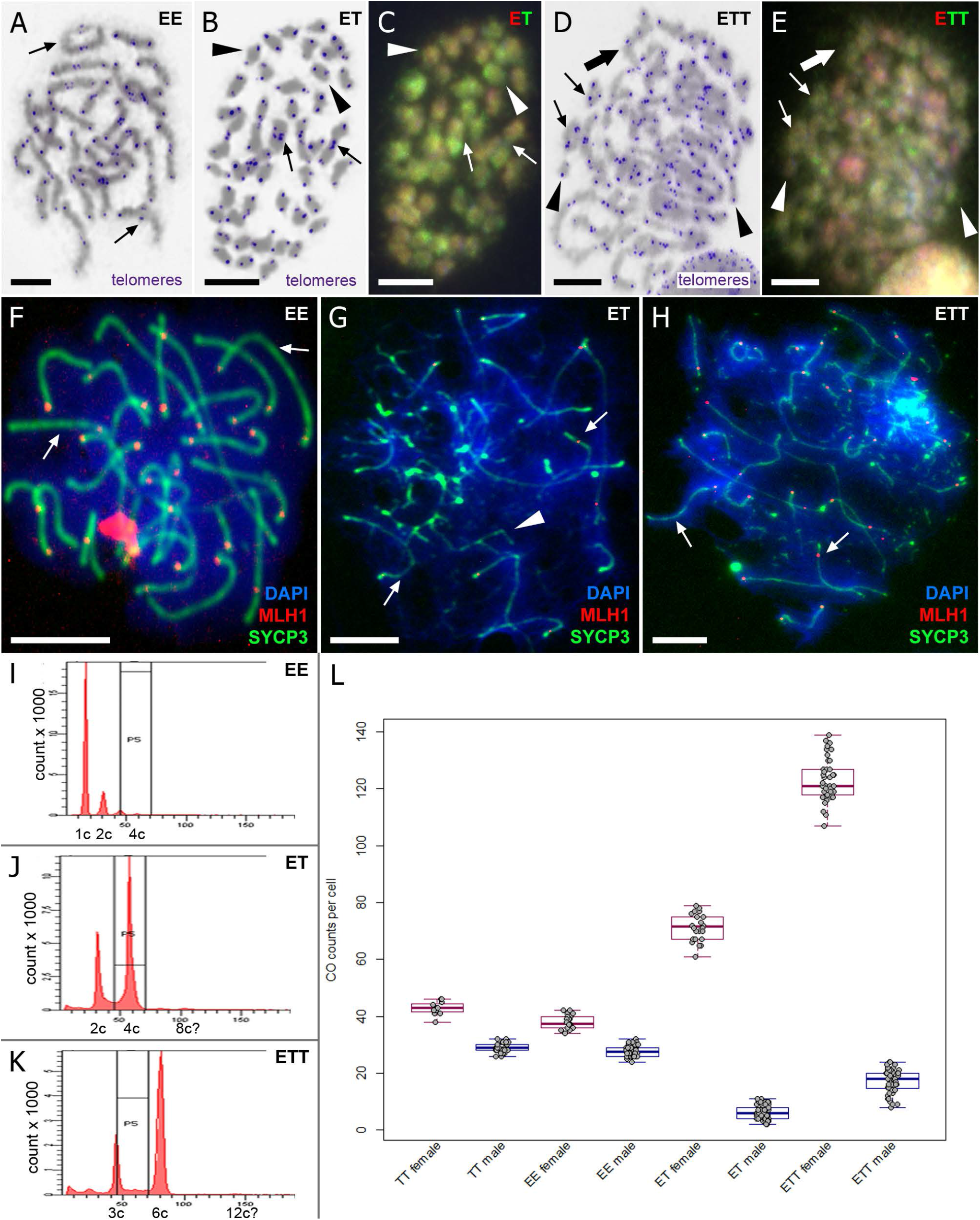
Male meiotic spreads at metaphase I (A-E) and pachytene (F-H) stages. Panels A, B, D show Giemsa stained chromosomes (grey) with FISH-labelled telomeres (blue) in *C. elongatoides*, ET and ETT hybrid males, respectively. Panels C, E show the same metaphases as in B and D, respectively, of hybrids but stained with CGH to reveal the origin of individual chromosomes (red dye corresponds to *C. elongatoides*-like chromosomes and green to *C. taenia*-like ones). Thin arrows indicate exemplary cases of bivalents, arrowheads exemplary univalents and thick arrows exemplary case of multivalent. Panels F-H show spread pachytene spermatocytes of *C. elongatoides* (F), hybrid diploid ET (G) and triploid ETT (H) males stained with DAPI (blue) in which syneptonemal complexes (SCs) were immunolabelled with antibodies against SYCP3 protein (green) and MLH1 protein (red). Arrows indicate exemplary bivalents, arrowheads show examples of abnormal pairing and failure of bivalent formation. Scale bar = 10µm. Panels I-K show the flow cytometry results of testis of *C. elongatoides* (I), hybrid diploid ET (J) and triploid ETT (K) males. Diagram in panel L shows the average frequencies of crossovers (COs) per cell in studied genomotypes (males and females indicated in blue and red, respectively).

Flow cytometric analysis also showed that hybrid males lacked the peak of 1C cells corresponding to haploid sperm nuclei characteristic for sexual males Fig 4I, corroborating the aforementioned data. By contrast, ET hybrids males possessed prominent peaks corresponding to diploid (2C) and double-diploid (4C) cell populations in gonads (this situation was analogous in ETT males with prominent triploid (3C) and double triploid (6C) cell populations) (Fig 4J, K). These 2C and 3C cell populations in ET and ETT males, respectively, correspond to spermatogonia and somatic cells while 4C and 6C cells, respectively, probably correspond to primary spermatocytes accumulated due to problems during chromosomal pairing. It is noteworthy that flow cytometry of hybrid male gonads revealed a minor cell population (less than 3%) with 8C and 12C DNA content in diploid and triploid hybrids, respectively (Fig 4J, K).

### Chromosomal pairing is restored in hybrid females but compromised in males

Low condensation of female diplotene chromosomes (lampbrush chromosomes) prevented the application of comparative genomic hybridization (CGH). Nevertheless, we identified the ancestry of particular chromosomes through bivalent morphology. We found that each parental species possesses several bivalents that may be used as species-specific markers due to unique distribution of characteristic structures (loops with specific morphology, spheres and nucleoli loci). In total, we recognized 9 marker-bivalents typical for *C. elongatoides* and 8 for *C. taenia*; S3 Fig upper panel). All these marker bivalents were present in hybrid females; diploid ET genomotype contained all 9 *C. elongatoides*-specific bivalents and 8 *C. taenia*-specific ones, while triploid ETT contained 9 *C. elongatoides*-specific bivalents and 16 *C. taenia*-specific bivalents, S3 Fig lower panel). The similarity of bivalents in hybrid females to those appearing in parental species suggests that they are formed by homologous chromosomes coming from the same species (i.e. ExE and TxT bivalents, respectively). Throughout this study, we will use the terms ‘homologs’ to define homologous chromosomes arisen from the same species while ‘orthologs’ to define homologous chromosomes arisen from different species. The appearance of aforementioned bivalents in hybrid females leads us to propose they are formed by identical copies arisen from endoreplication.

In contrast to their hybrid sisters, metaphase nuclei of diploid hybrid males exhibited many aberrant structures. The successful application of CGH to male figures demonstrated that bivalents and multivalents in ET males contained pairing chromosomes derived from different parental species (Fig 4C). We also noticed that both genomes contributed roughly equally to the formation of univalents (i.e. on average we observed ~ 16 T-like and ~ 18 E-like univalents). Metaphases of triploid ETT males possessed a higher proportion of bivalents that were formed by TxT chromosome pairing (probably homologues), but several instances of ExT chromosomal pairing were also observed. Some multivalents appeared to contain chromosomes of both species, and univalents were usually composed of E-like chromosomes but T-like univalents were occasionally observed (Fig 4E).

### Chromosomal pairing aberrations are initiated during early stage of bivalent formation

Immunostaining for SYCP3 proteins showed well developed lateral elements of synaptonemal complexes (SC) of 24 and 25 bivalents during the pachytene-stage nuclei of *C. elongatoides* and *C. taenia* males and females, respectively (Fig 3E, Fig 4F, S5 Fig A). Additionally bivalent formation was confirmed by immunostaining with SYCP1 (central component of SC). Similarly, clonal ET and ETT females had well developed SCs along the entire lengths of their 49 and 73 bivalents, which corresponded to aforementioned observations during the diplotene stage (Fig 3F, G, S5 Fig B). By contrast, hybrid ET males generally contained only short and partially formed SCs, often branching and forming loops, indicating improper pairing of chromosomes. Rarely, we observed one or two normal looking SCs in these males (Fig 4G, S4 FigA, S5 Fig C). In triploid ETT males we also observed many malformed SCs but, contrary to ET diploids, between 8-15 properly developed SCs were usually present per cell (S5 Fig D). This corroborates the hypothesis that certain TxT bivalents form properly in ETT triploid (Fig 4H, S4 FigB). In abnormal bivalents of diploid and triploid hybrids the SYCP3 was localized to subtelomeric regions while inner fragments of chromosomes usually lacked the SYCP3 signals (S4 Fig).

### Frequency of crossovers differs between sexes and species and is decreased in hybrids, especially in males

The occurrence and rate of meiotic crossovers (COs) could be traced in diplotene metaphase I stage nuclei as chiasmata or in mid- and late pachytene nuclei as antibody staining locations of MLH1 sites (DNA mismatch repair protein) [25,26]. Depending on the stage of investigated cells, we scored either one or the other data type. We note that in most females both chiasmata and MLH1 sites were available and we confirmed they provide equivalent counts of CO sites. Investigated individuals of both parental species always contained one or more chiasma or MLH1 focus per bivalent indicating that crossovers always take place between paired chromosomes. However, we found significant differences in CO frequencies between species with generally higher values in *C. taenia*, and also between sexes. Similar to *Danio rerio* [27], the CO sites of males were less numerous and mostly located close to the telomeric regions of bivalents, while female COs were more numerous and rather evenly distributed. On average, *C. taenia* males and females had ~28.9 and ~42.8 COs per cell, respectively and *C. elongatoides* males and females had ~27.7 and ~38 COs per cell, respectively, (Fig 4L). The mixed effect of the Generalized Linear Model GLM_poisson_ incorporating both parameters fitted the data significantly better than single parameter models (formula = CO~sex+species+(1|individual_ID); p(test=ChiSq) = 0.04).

Female hybrids also expressed one or more chiasma or MLH1 focus per bivalent, but the CO frequency in the ET genomotype was slightly but significantly lower than predicted by simple extrapolation of sexual counts to endoreplicated hybrid nuclei; we observed only ~71.3 COs against ~80.8 expected (one sample t-test p < 10^−5). ETT females had ~123.4 COs per cell, which almost exactly fit the expectation value (123.6). The decrease in CO frequencies caused by hybridization was much more drastic in males, which possessed only ~6.2 (diploid ET) and ~17.2 (triploid ETT) MLH1 foci per cell (Fig 4L). Although the CO frequency was partially increased in triploids, it was still much lower than expected.

## Discussion

Mechanisms underlying the emergence of reproductive isolation barriers, and hence the existence of biological species have been intensively studied since Darwin’s time, as has the question of why sex is a dominant reproductive mechanism among Metazoans. Our study shows that to a surprisingly high extent, both questions are complementary. *C. elongatoides* and *C. taenia* diverged approximately 9 Mya and despite intensive historical introgressions, their speciation is now complete due to the absence of contemporary gene exchange [9]. In this study we identified gametogenetic aberrations, which exist in both male and female hybrids and represent efficient postzygotic barriers to gene flow. However, the cytogenetic background of such barriers radically differs between hybrid sexes (Fig 1 B-C); while females produce clonal gametes via premeiotic endoreplication, males most likely lack this mechanism and are sterile. Flow cytometry indicated minor populations of octoploid or dodecaploid testicular cells in ET and ETT males, respectively (Fig 4J, K), which may be theoretically interpreted as a result of premeiotic endoreplication of spermatocytes [28]. However, given that we never observed any testicular cells with double the number of chromosomes despite cytogenetic analysis of hundreds of metaphase I nuclei, such a hypothesis seems unlikely. The occurrence of unusually high-ploidy cells may indicate meiotic arrest after chromosome replication [29] or, given larger heads and multiple axonemes or flagella in rarely occurring hybrid spermatozoa, these cells may represent fused or unseparated spermatozoa.

In any case, while premeiotic endoreplication is a dominant gametogenetic pathway in hybrid females, it seems absent or at least very rare in males where the major aberration associated with their sterility is the failure of chromosomal pairing and/or reduction of crossovers, which prevent proper segregation of chromosomes. These aberrations likely enforce the meiotic arrest at the pachytene-metaphase I followed by apoptosis in the majority of spermatocytes. Because hybrid males suffer from improper formation of SCs and drastically decreased CO rates, we propose that these aberrations take place during the early stages of homologue pairing. Such stages are characterized by preliminary Double Strand Break (DSB)-independent pairing that restricts the searching area for homologous recognition and subsequent DSB formation and co-alignment [30]. In cypriniform fishes, the homology search is initiated in subtelomeric regions, followed by SYCP3 upload and zipping towards the interstitial chromosomal segments [31]. In case of sufficient homology between two chromosomes, zipping is finalized by uploading of SYCP1 protein accomplishing bivalent formation [31]. It is therefore possible that low homology between orthologous chromosomes of contemporary species prevented sufficient pre-DSB coalignment in their hybrids, which instead possessed ectopic recombinational intermediates between similar sites of nonhomologous chromosomes. Such ectopic complexes are supposedly less stable and probably eliminated by mismatch repair systems thereby preventing their maturation to MLH1 sites [14], which leads to SC formation around the initial subtelomeric segments and generally a low CO frequency.

Gametogenetic aberrations in hybrids may take various forms spanning from mitotic arrest in germ cells before meiosis [32] to sperm production without meiotic division [33], but several studies reported patterns analogous to the *Cobitis* case [14,18,26], suggesting that compromised pairing and reduced CO frequency can be a relatively common aberration underlying HS. The proximate molecular mechanisms underlying HS remain generally unknown, but researchers have traditionally preferred genic models (i.e. Dobzhansky-Muller type incompatibilities) over the role of divergence in DNA base composition (the Bateson’s model). Nevertheless, recent progress suggests both aspects could operate together since sterility of mouse hybrid males probably results from divergence between the PRDM9 gene responsible for the positioning of recombination sites and its recognition sites resulting in asymmetrical DSBs and gametogenetic abruption [34]. The importance of sequence homology in HS was further underlined by evidence that fertility of mouse hybrid males drastically increased when homology was restored to segments of chromosomes that were mainly involved in mispairing [34]. Observations of *Cobitis* point to the same conclusion since ETT triploids, albeit sterile, had partially rescued bivalent formation and increased CO rate due to proper pairing of *C. taenia*-like homologues. Similarly, in *Darevskia* lizards the CO rate of triploid male hybrids matched the values in parental species contributing double chromosome sets [18]. These findings not only support the importance of sequence homology in hybrid gametogenesis but they also indicate that the remedy of HS may, at least partially, be delivered by polyploidization, which adds extra copies of genomes conspecific to one or both parents. In this context it is indeed noteworthy that many hybrid animals and plants are polyploids and form bivalents from conspecific homologues [35,36].

Our findings indicate that asexuality may represent a surprisingly efficient remedy to hybrid sterility that arises during species differentiation. Indeed, if sterility is largely caused by the problem of orthologous chromosome pairing, then premeiotic endoreplication observed in various asexual taxa could provide an elegant solution by supplying each chromosome a peer to properly pair with. Nevertheless, the reality appears more complex since we observed different pairing affinities of homologues and orthologues in males and females. In particular, while triploid ETT males always possessed some T-like univalents and ExT bivalents, even though each *taenia*-derived chromosome had a conspecific homologue, hybrid females always formed perfect sets of ExE and TxT bivalents without any observed mismatches. Improper pairing of homo vs. orthologues is likely detrimental and consequently, for example, neopolyploid lineages tend to shift from polysomic to disomic inheritance in order to gain stability [3]. In some instances the avoidance of multivalents and orthologous pairing might have been achieved by increased CO interference causing reduction of CO rate between orthologs[3]. On the other hand mispaired chromosomes can lead to compensatory mechanism increasing COs in paired ones suggesting genic regulation of CO frequency [37]. Recent evidence also indicated that activity of specific genes may selectively prefer homologous over orthologous pairing [38]. Our findings that orthologous pairings and significantly decreased CO rate occur in polyploid males spermatocytes but not in females oocytes having only endoreplicated homologs paired and similar relative CO rate to sexual species indicates the existence of some basic differences between sexes in mechanisms of chromosomal pairing or homology searching. For example, females have generally more CO sites per chromosome, which are localized not only to telomeric regions but also to interstitial segments. This may increase the incidence of high-homology interactions and hence, unlike their brothers, females would potentially have a more stringent disruption of low homology or ectopic pairings, ultimately ensuring successful formation of bivalents from homologues and avoidance of orthologous pairing. The process of homology search during early meiotic stages that ensures the proper pairing of chromosomes is not well understood especially in animals and represents a challenge in current molecular and cellular biology [3]. Testing whether pairing of homologous vs. orthologous chromosomes differ so strikingly between males and females also in other asexual complexes and revealing the reasons why, may uncover some basic rules of this process.

In any case, the fact that endoreplication prevails in females but not in males is a striking outcome of the present study. In speciation research the asymmetrical accumulation of gametogenetic aberrations is a well-known and intensively studied phenomenon (e.g. Haldane’s rule) and Torgasheva and Borodin [14] recently proposed that during species differentiation both hybrid sexes may follow the same route to sterility caused by incorrect homologous pairing and aberrant synapses, just at a different pace. The analysis of the *Cobitis* asexual complex somewhat challenges this view as it shows that even when postzygotic barriers have been accomplished in both hybrid sexes, the underlying mechanisms may have very different backgrounds because hybrid females remained fertile but restricted gene flow by the production of clonal unreduced gametes. Premeiotic endoreplication is a relatively common pathway among asexual plant and animal hybrids (rev. in [2,39]) but to date its triggers have not been revealed, which arguably represents a considerable gap in holistic understanding of speciation and the evolution of sex. The present investigation of *Cobitis* puts the occurrence of hybrid asexuality into a context with hybrid sterility and shows that it likely arises as a sex-specific process. Theoretically, one may hypothesize that endoreplication is equally rare in both sexes upon initial hybridization but since successful clones may be established only by successfully reproducing females, natural selection would favour females with high rates of endoreplication, thereby making the impression of sex-specific differences. However, Choleva et al. [21] reported that clonality is pervasive already among F1 elongatoides-taenia females, making this explanation unlikely.

So, what causes endoreplication and why is it specific to females? Although we, just like any other study to date, did not witness the particular moment of endoreplication nor did we reveal its causal stimulus, comparison of our data with the related genus *Misgurnus* leads us to propose that it is not determined by hormonal levelsbut more likely by genetic sex determination. Specifically, Yoshikawa et al. [28] artificially reversed the sex of diploid hybrid *Misgurnus* females into males and reported the production of unreduced spermatozoa via endoreplication, suggesting that diploid individuals genetically determined as females possess the endoreplication ability regardless of their gonadal phenotype. The sex-determination system in the *C. taenia* complex is unknown, but male heterogamety (X_1_X_2_Y or X0, respectively) has been reported in congeneric species [40,41], making it tempting to speculate that intersexual differences in fertility and asexuality relate to the sex determination system. However, if this is the case, we note that sterility of putatively heterogamic males would conform to the famous Haldane’s rule only superficially, since the fertility of females is ensured by the premeiotic endoreplication that is absent in males. Although females should theoretically face the same complications of low homology between chromosomes from both species, their extraordinary gametogenetic modification would circumstantially complement HS by pairing the endoreplicated sister chromosomes and ultimately allowing clonality.

Morishima et al. [42] reported that artificially induced tetraploid *Misgurnus* hybrids lost the capacity of endoreplication present in their diploid hybrid ancestors, suggesting that the ability of endoreplication somehow depended on ploidy and may be under dosage-dependent control of some crucial transcripts. Indeed, the emergence and stability of asexual lineages is known to depend on particular ploidy levels [5] and reproductive strategies of hybrids between the same species may drastically differ, depending on whether they are diploid or polyploid [42–45]. However, the *Cobitis* case shows that the effects of ploidy are not so strict since both diploid and triploid *C. elongatoides-taenia* females have similar gametogenetic pathways and may establish very successful clonal lineages in nature [46].

It was traditionally hypothesized that induction of asexuality in hybrids is possible upon combinations of only certain species that possess some specific predispositions to it [47,48]. In contrast to this view, our study documented that asexuality arises relatively frequently upon genome merging of various sexual species. Namely, we found the same type of gametogenetic mechanism in *C. elongatoides-taenia* hybrid females as that reported in distantly related *Misgurnus* hybrids from Asia [28,49], making it unlikely these two distant lineages share the same predisposition. Moreover, we proved by comparative analysis that in fishes that asexual hybrids tend to arise between distantly, rather than closely related species [9] and demonstrated that while *C. elongatoides* and *C. taenia* produced fertile non-clonal hybrids during the early stages of their diversification, only asexual and sterile hybrids were formed after ongoing divergence.

How genetic divergence between species may mechanistically trigger asexuality of their hybrids is unknown, but it is interesting that several previously proposed explanations are conceptually similar to either genic or non-genic views of speciation. Namely, the Balance Hypothesis [5] matches the Dobzhansky-Muller genic models by postulating the prominent role of gene-to-gene interactions, De Storme & Mason’s model is rather similar to [50]’s view of non-genic residues, as implemented in chromosomal speciation models [51], and Carman [7] envisaged the roles of diverged regulatory networks in the establishment of hybrid asexuality, which is similar to Tulchinsky et al.’s [52]’s view of postzygotic trans-regulatory incompatibilities. Thus, for example the reason why the majority of known asexual hybrids are formed from parental species with a certain level of divergence may be that key genetic pathways affected by accumulated incompatibilities fail to suppress the replication of hybrid chromosomes before cytokinesis, ultimately resulting in endoreplication. However an alternative explanation may assume that increased divergence is not needed as a trigger for asexuality, but rather may prevent pairing of orthologous chromosomes. In other words, endoreplication may appear also in hybrids between close species but decreased homology of orthologs is necessary to prevent the formation of spurious bivalents or multivalents after endoreplication. Research on asexual organisms has already contributed to many domains of biology but their gametogenetic pathways are rarely understood. This study shows there is a tight link between hybrid asexuality and sterility leading us to believe that thorough investigation of gametogenetic aberrations in these interesting organisms would considerably increase not only our understanding of speciation, but also of meiosis and chromosomal pairing.

## Methods

### Samples studied

The identity, ploidy and genome composition of every investigated specimen was evaluated with standard species-diagnostic markers [46]. Sex of each individual was a priori as for males, we in total investigated 3 *C. taenia* (TT), 3 *C. elongatoides* (EE), 5 diploid hybrid (ET) and 4 triploid hybrid (ETT) individuals. As for females, we in total analyzed 4 TT, 4 EE, 3 diploid hybrid ET and 5 triploid hybrid ETT individuals. All females and the triploid ETT males came from natural sites, while diploid ET males represent F1 generation from experimental breeds. This is because hybrid males do not form self-perpetuating lineages and hence in nature only triploid forms are occasionally found as results of ET females backcrossing to parental species, while true diploid F1 male generation has never been sampled in natural sites.

### Histology and SEM

Small fragments of the testes for light and electron microscopy were fixed at 4 °C in 2.5% glutaraldehyde with phosphate buffer at ph 7.4 and post-fixed in 1% osmium tetroxide using the same buffer. After dehydration with ethanol series, the tissues were embedded in Epon 812. Semi-thin sections were stained with methylene blue while ultrathin sections of the selected areas were contrasted with uranyl acetate and lead citrate and analyzed using a JOELJEM-100Sx transmission electron microscope. Apoptotic cells were detected with the QIA33 | FragEL™ DNA Fragmentation Detection Kit

### DNA flow cytometry

Genome size of cell populations from the testes was estimated by measurement of cells nuclei using BD FACSAria™ II flow cytometer. Detail description of methods is given in S1 Methods.

### Mitotic and meiotic metaphase chromosomes

Mitotic and meiotic metaphase chromosomes spreads were obtained from kidneys and testes of sexual and hybrid males without colchicines injection according to standard procedures [53]. Metaphases chromosomes were initially stained with Giemsa in order to check the number and morphology of chromosomes and bivalents formation.

### Pachytene chromosomes and immunofluorescent staining

Pachytene chromosomes spreads were done from testes according to the protocol described in [54] and from ovaries following [55] protocol. Synaptonemal complexes (SCs) were visualized using immunofluorescent staining with antibodies (atb) against SYCP3 (ab14206, Abcam) and SYCP1 (a gift from Sean M. Burgess). The recombination sites were visualised by atb staining against the MLH1 (ab15093, Abcam) proteins. Detail description of pachytene chromosome preparation and immunofluorescent staining is given in S1 Methods.

### Fluorescent *in situ* hybridization and Comparative genomic hybridization

FISH with a commercial PNA probe for TTAGGG sequence was performed on metaphase slides and slides with male pachytene chromosomes according to the protocol provided by manufacturer (DAKO). Comparative genome hybridization (CGH) was made according to [56] on the same slides where telomeric sites were detected by FISH. Detail description of CGH is given in S1 Methods.

### Diplotene chromosomes

Chromosomes during diplotene stage (lampbrush chromosomes) were microsurgically isolated from growing oocytes of parental and hybrid females according to the protocols developed for amphibians [57] with modifications suggested by [58]. Description of bivalent morphology and lampbrush chromosome maps construction were performed according to Callan’s monograph in Corel™ DRAW graphics suite X8 software. Detail description of diplotene chromosome preparation is given in S1 Methods.

## Acknowledgements

Authors are obliged to Roger Butlin, František Marec, Lukáš Kratochvíl and Radka Reifová for inspiring comments on the Manuscript and thank to Lukáš Choleva for help with fish maintenance, to Karel Halačka for his assistance in the field and to Sean M. Burgess for sharing antibodies against SYCP1. We are grateful for financial support from the Czech Academy of Sciences grant no. RVO67985904, Czech Science Foundation grant nos. 17-09807S and 19-21552S, the Ministry of Education, Youth and Sports of the Czech Republic grant EXCELLENCE CZ.02.1.01/0.0/0.0/15_003/0000460 OP RDE and National Science Centre, Poland: grants no. DEC-2011/03/B/NZ8/02095 (to JK) and 2011/03/B/NZ8/02982 (to AB & DJ). The funders had no role in study design, data collection and analysis, decision to publish, or preparation of the manuscript.

## Author contribution

DD, ZM and KJ designed the study and co-drafted MS. DD, ZM and AM performed cytogenetic analyses. DJ, AB, JKo provided material for experiments. JKl, DD and AM performed flow cytometry. MP performed histological experiments, transmission electron microscopy and wrote relevant parts of the text. KJ performed statistical analyses and drafted first version of the text. All co-authors contributed to final text version.

