## Supplemental Methods for "Parthenogenesis as a solution to hybrid sterility: the mechanistic basis of meiotic distortions in clonal and sterile hybrids"

**Supplementary Methods**

**DNA flow cytometry:** Testis were homogenized manually followed by 0.15% trypsin treatment to separate individual cells. Blood samples and sperm suspension of sexual species were used as internal control. Testis cell suspension and blood samples were incubating with 0.1% Triton X100, 10 μ/ml DAPI and 15 mM $MgCl_2$ for 4–6 h (at +4°C) to realize cell nuclei which were subsequently measured by BD FACSAria™ flow cytometer. At least 10,000 events were measured. Data were further analyzed by BD FACSDiva software (version 6.1.3).

**Pachytene chromosomes and immunofluorescent staining:** Pachytene chromosomes were obtained for males and females according to protocols described in [1,2]. After manual homogenization of male gonads, 1 µl of suspension was placed in the 30 µl of hypothonic solution (1/3 of 1× PBS) preliminary dropped on SuperFrost® slides (Menzel Gläser) for 20 minutes and subsequently fixed in 2% paraformaldehyde for 4 minutes. After fixation slides were washed 1× PBS and used for immunofluorescent staining (IF). In case of female, 20 µl of manually homogenized cells suspension from ovaries was put on SuperFrost® slides (Menzel Gläser) followed by addition of 40 µl of 0.2 M Sucrose and 40 µl of 0.2% Triron X100 for 7 min and subsequently fixed in 2% PFA for 16 minutes. After washing in 1x PBS slides were stored until IF procedures.

Synaptonemal complexes (SC) of chromosomes during pachytene were visualized using immunofluorescent staining (IF) with rabbit polyclonal antibodies (ab14206, Abcam) against SYCP3 protein which is the lateral component of SC and chicken polyclonal SYCP1 (a gift from Sean M. Burgess) which is the central component of SC. Recombination foci were identified using mouse monoclonal antibodies (MLH1 (ab15093, Abcam) against MLH1 protein, mismatch repair protein. Fresh not dried slides were incubating with 1% blocking reagent (Roche) in 1x PBS and 0.01% Tween-20 for 20 min and then with primary antibody (dilutions as recommended by manufacturers) for 1h at RT. Slides were washed in 1x PBS at RT and incubated in combination of secondary antibodies (Cy3-conjugated goat anti-rabbit IgG (H+L) (Molecular Probes) and Alexa-488-conjugated goat antimouse IgG (H+L) (Molecular Probes) for 1h at RT. Slides were washed in 1x PBS with 0.05% Tween-20 and mounted in Vectashield/DAPI (1.5 mg/ml) (Vector, Burlingame, Calif., USA).

**Fluorescence *in situ* hybridization and Comparative genomic hybridization:** Fluorescence *in situ* hybridization (FISH) with probe to telomeric repeat was performed on metaphase slides and slides with pachytene chromosomes after IF staining in order to precisely distinguish bi-, uni- and multivalent. A commercial PNA probe to TTAGGG sequence was used for FISH according to protocol provided by manufacturer (DAKO).

To distinguish whether bivalents in hybrids are formed between homologous or orthologous chromosomes we performed comparative genome hybridization (CGH) on meiotic metaphase and lampbrush chromosomal spreads [3]. For CGH experiments whole genomic DNA (gDNA) of pure parental species *C. elongatoides* and *C. taenia* was used to prepare probes. gDNA was extracted from muscles or fins using the DNeasy Blood and Tissue Kit (Qiagen, Hilden, Germany) according to the manufacturer's instructions. *C. elongatoides* gDNA was labelled with biotin-16-dUTP (Roche, Mannheim, Germany); *C. taenia* gDNA was labeled with digoxigenin-11-dUTP (Roche, Mannheim, Germany) using a Nick Translation Mix (Abbott) following the protocol supplied by the manufacturer. The best results were obtained after 2.5 – 3 hours of nick translation until labeled DNA fragments were approximately 200–500 bp long. In case of meiotic metaphase chromosomes we performed CGH on the same slides where we detected telomeric sites. Probe of both parental species were mixed together and with hybridization mixture (50% formamide, 2x SSC, 10% dextran sulphate, salmon sperm DNA). Hybridization was performed at 37ºC for 48 hours, followed by post-hybridization washes with 50% formamide in 2× SSC at 42°C for 5 min (3 times) and 2× SSC for 5 min (3 times). The biotin-dUTP and digoxigenin-dUTP labelled probes were detected using streptavidin-Cy3 (Invitrogen, San Diego, Calif., USA) and antidigoxigenin-FITC (Invitrogen, San Diego, Calif., USA) correspondingly. The chromosomes were counterstained with Vectashield/DAPI (1.5 mg/ml) (Vector, Burlingame, Calif., USA).

**Diplotene chromosomes:** Diplotene chromosomal spreads (lampbrush chromosomes) were prepared from parental and hybrid females according to the protocol initially developed for amphibian oocytes [4] with modifications suggested in [5]. Vitellogenetic oocytes of 0.5–1.5 mm in diameter were taken from non-stimulated females within the OR2 saline (82.5 mM NaCl, 2.5 mM KCl, 1 mM $MgCl_2$, 1 mM $CaCl_2$,1mM $Na_2HPO_4$, 5 mM HEPES (4-(2-hydroxyethyl)-1-piperazineethanesulfonic acid); pH 7.4). Nuclei were then microsurgically isolated from oocytes by jeweler forceps and needles in the isolation medium "5:1" (83 mM KCl, 17 mM NaCl, 6.5 mM $Na_2HPO_4$, 3.5 mM $KH_2PO_4$, 1mM $MgCl_2$, 1 mM DTT (dithiothreitol); pH 7.0–7.2). Nuclear envelopes were manually removed in one-fourth strength "5:1" medium with the addition of 0.1% paraformaldehyde and 0.01% 1M $MgCl_2$ in a chambers attached to a slide meaning that in each chamber we obtained chromosome spread from individual oocytes. Slide with oocyte nuclei contents were subsequently centrifuged for 20 min at +4°C, 4000 rpm, fixed for 30 min in 2% paraformaldehyde in 1x PBS, and post-fixed in 70% ethanol overnight (at +4°C). Description of bivalents morphology and lampbrush chromosome maps construction were performed according to Callan [4] in Corel™ DRAW graphics suite X8 software.

**Wide-field and fluorescence microscopy:** Mitotic and meiotic chromosomes after FISH, CGH and IF were inspected using Carl Zeiss Axio Imager.Z2 and Provis AX70 Olympus microscopes equipped with standard fluorescence filter sets. Microphotographs of chromosomes were captured by CCD camera (DP30W Olympus) using Olympus Acquisition Software and CoolCube 1 using Metasystem platform for automatic search, capture and image processing. Microphotographs were finally adjusted and arranged in Adobe Photoshop, CS6 software.
