## Supplementary figures and images for "Parthenogenesis as a solution to hybrid sterility: the mechanistic basis of meiotic distortions in clonal and sterile hybrids"

### Supplemental Figures

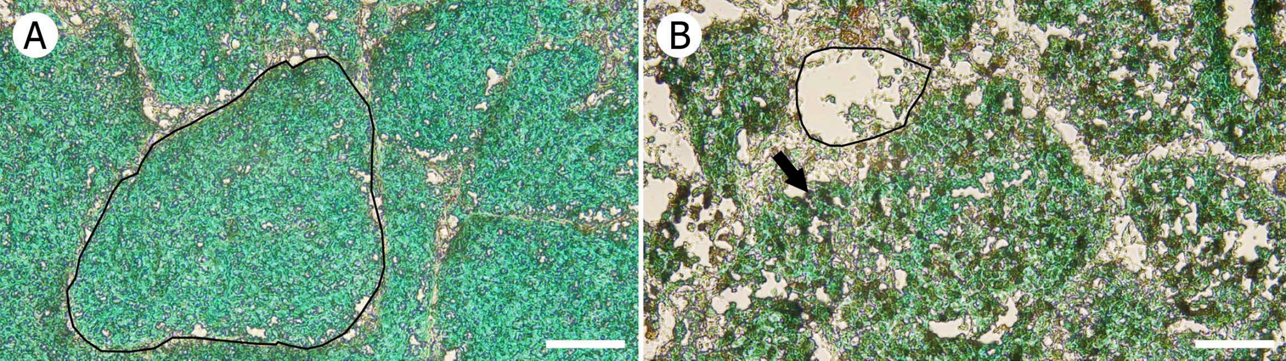
